# Kinetic isotope tracing of glycerol and de novo proteogenic amino acids in Human Lung Carcinoma cells using [U-^13^C_6_]glucose

**DOI:** 10.1101/2020.10.28.358432

**Authors:** Subia Akram, Jyotika Thakur, Manu Shree, Shyam K. Masakapalli, Ranjan Kumar Nanda

**Author notes:** **Correspondence: Ranjan Kumar Nanda (PhD)**, Translational Health Group, International Centre for Genetic Engineering and Biotechnology, New Delhi – 110067, E. mail.

## Abstract

13C based Isotopic tracers of the media components can be used to kinetically track their contribution in the cell systems. A tracer ([U-^13^C_6_] glucose) was used to monitor its contribution into the central carbon metabolic pathways of human lung carcinoma (A549) cells by Gas chromatography-mass spectrometry (GC-MS) based mass isotopomer analysis. Calculated average 13C of methanolic extracts (glycerol: 5.46±3.53 % and lactate: 74.4±2.65 %), protein acid hydrolysates (serine: 4.51±0.21 %, glycine: 2.44±0.31 %, alanine: 24.56±0.59 %, glutamate: 8.81±0.85 %, proline: 6.96±0.53 % and aspartate: 10.72±0.95 %) and the culture filtrate (glycerol: 43.14±1.45 % and lactate: 81.67±0.91 %), showed significant contribution of 13C glucose. We observed the Warburg effect with higher levels of 13C lactate in the culture filtrate. 13C glycerol levels in culture supernatant showed significant increase with time and amino acids of glucogenic origin also contributed cellular protein biomass. The workflow adopted in this study for 13C analysis could be useful for the metabolic phenotyping of other mammalian cell systems under normal and perturbed (cancer and infection) conditions.

## 1. Introduction

Nutrient availability is vital in maintaining a local immune environment in tumour or infected tissues for effective host immune response. Tumour cells and pathogens compete with host immune cells for nutrients as a part of their immune evasion strategies. Resulting nutrition imbalance leads to impaired immune function and compromised outcome in immuno-therapies [1]. Upon infection or introduction of chemical signals, cancer cells and activated immune cells adopt metabolic reprogramming to acquire biomass for extensive proliferation by coordinating systemic oxidation of carbon, nitrogen and other nutrients [2,1]. Understanding the altered cellular metabolism to meet the biomass demand, kinetically, will allow identification of important pathways for targeted, effective therapeutic interventions.

The duplication rate of cancerous and immune cells varies widely (17.0-79.5 hours) based on their origin and types [3]. Moreover, proteins contribute to ~17-26% of the biomass, and its synthesis rate influences the cell duplication period. However, glucose or glutamine and its intermediates contribute <10% of cell mass [4]. Like nucleotides, after synthesis of sufficient protein pool, the cells become ready for duplication. Protein synthesis in immortalised and primary cells in culture depends on amino acids present in culture media and de-novo amino acid synthesis. In non-dividing mammalian cells, the ratio of externally supplied versus endogenous amino acids to de novo protein synthesis was about 17:1 [5].

While, 13C tracers analysis have been a promising tool to define cellular and organ-specific central metabolism (glycolysis, tri-carboxylic acid, pentose phosphate pathways), additional studies to quantify the contribution of nutrients to de novo amino acid synthesis directed towards protein synthesis are essential [6,7,8]. Elucidation of the source of amino acids and its incorporation in new protein synthesis will be informative to identify essential steps in cell metabolism to develop interventions for different conditions like cancer and infectious diseases. This study quantified the contribution of amino acids originated from glucose metabolism and used in protein biosynthesis of A549 cells kinetically supporting the cancerous metabolic phenotype.

## 2. Results

### 2.1 Experimental workflow/

Type-II alveolar epithelial (A549) cells were grown in the presence of [^13^C_6_]glucose (99.6 %, 17.5 mM). Metabolites isolated from the cell culture filtrate, methanolic cell extracts and acid hydrolysate of protein fractions were analysed using gas chromatography and mass spectrometry (GC-MS) after derivatisation (Figure 1A). A549 cells grew linearly and reached a pseudo isotopic steady-state metabolism by 48 hours, ideal for 13C based metabolic flux analysis (13C-MFA) (Figure 1B).

**Figure 1.**
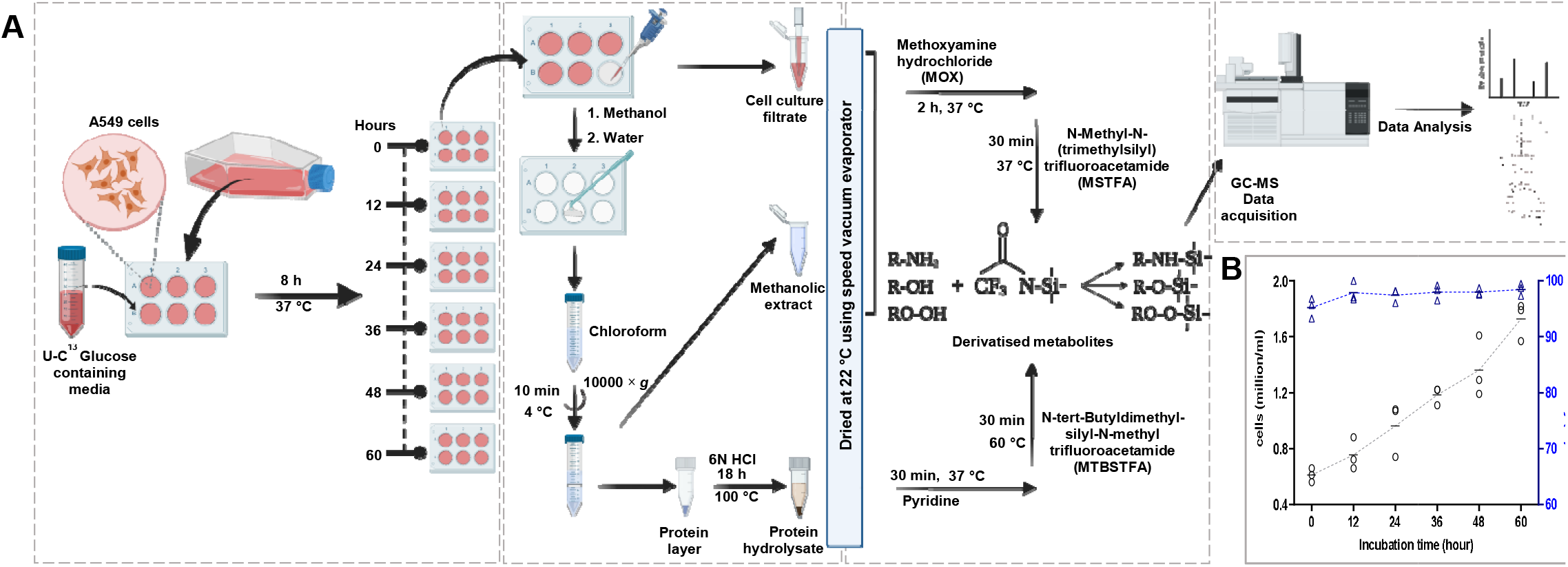
Kinetic isotopic labelling of A549 cell metabolites with [U-13C] glucose. (A) Workflow design of 13C labelling experiment. (B) Growth kinetics and viability of A549 cells cultured in vitro.

#### 2.2 13C Mass isotopomer incorporation

Efflux of labelled 3C units (glycerol: 43.14±1.45 % and lactate: 81.67±0.91 %) in normoxic A549 cells were observed (Figure 2A, 2B). A marked glycerogenic flow provided glycerol (5.46±3.53 %) as a final 3C export substrate together with lactate (74.4±2.65 %) (Figure 2C). To an extent, there is de novo synthesis of Serine (4.51±0.21 %) and Glycine (2.44±0.31 %) with a significant contribution from the media. (Figure 2D). Lactate (74.4±2.65 %), Alanine (24.56±0.59 %), Glutamate (8.81±0.85 %), Proline (6.96±0.53 %) and Aspartate (10.72±0.95 %) were also observed to have 13C labelling (Figure 2D).

**Figure 2.**
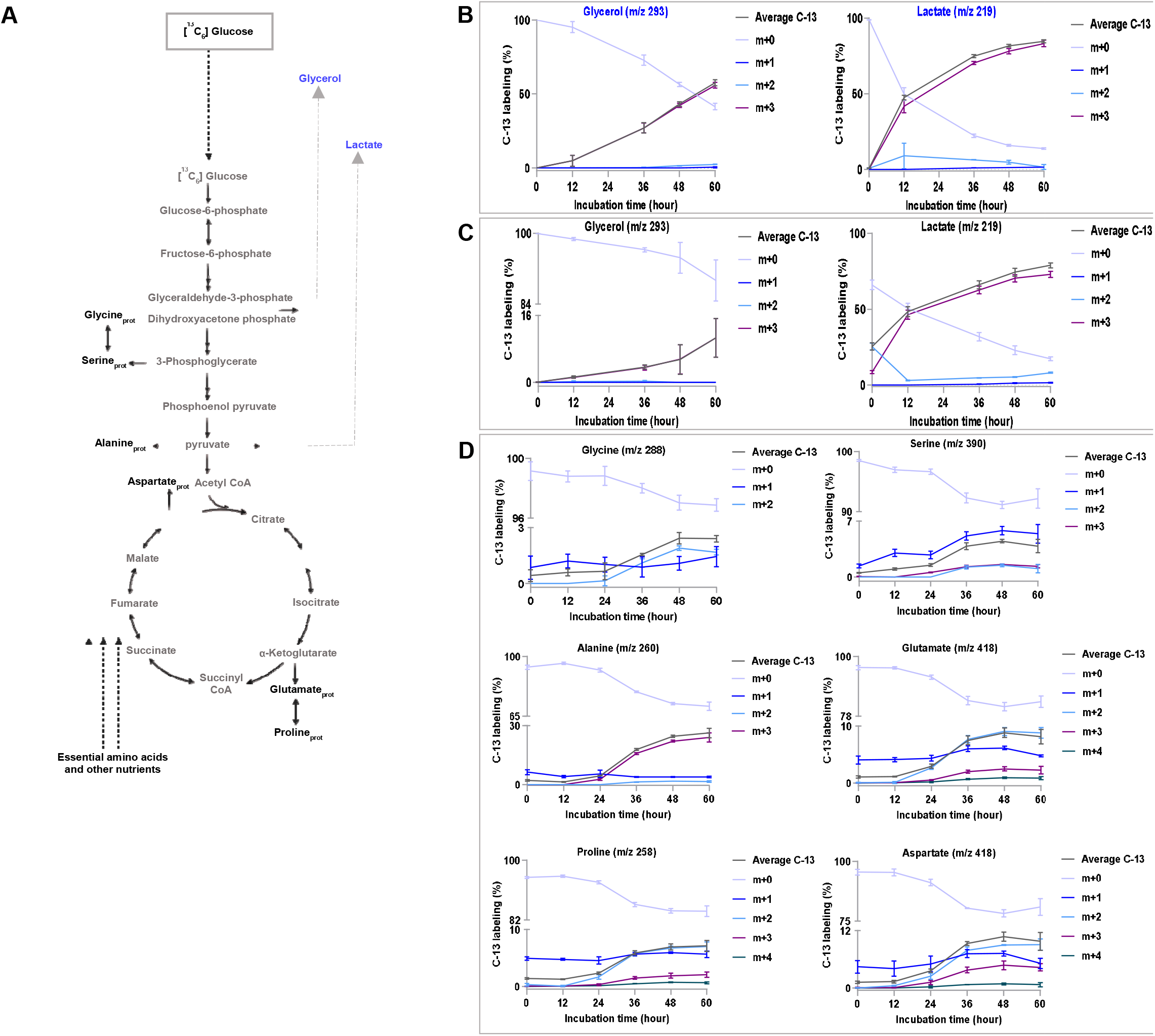
Mass isotopomer analysis. (A) Incorporation of 13C from [^13^C_6_] glucose through glycolysis and Krebs cycle. (B, C and D) Average 13C and individual mass isotopomer distributions (MIDs) in % of metabolites from cell culture filtrate, methanolic extract and acid hydrolysate protein (Amino acid_pro_) samples. The MIDs were corrected for natural isotope abundances. Data represents n = 3 replicates ± SD.

## 3. Discussion

Lactate and glycerol production in cells is assumed to contribute significantly to convert glucose to 3C units, thus lowering the adverse effects of excess glucose. Lactate and glycerol also act a signalling molecules and provide the metabolic phenotype of cancer and immune cells by regulating multiple gene expressions [9]. De novo synthesis of serine and glycine might support the 1-Carbon metabolism needed for cell proliferation. Pyruvate from C13 glucose contributes to labelled lactate and alanine. Subsequent, mitochondria metabolism lead to the synthesis of labelled glutamate, proline and aspartate. These amino acids are utilised in metabolic pathways and also get incorporated into the synthesis of new proteins. Essential amino acids (threonine, lysine, valine, tyrosine, phenylalanine) in the protein hydrolysate were unlabelled and might be contributed from the culture media. GC-MS based mass isotopomer analysis of the metabolites from the culture filtrate, methanolic fraction, amino acids derived from acid hydrolysates of protein allowed us to capture: i) the relative contribution of glucose vs other carbon sources on metabolism, ii) the rate of C13 incorporation in amino acids shedding light on potential pathway activities, by employing an, iii) optimised analytical conditions concerning ideal derivatising reagents and GC-MS parameters.

One of the limitations is that the contribution of C13 glucose generated through gluconeogenesis was not quantified in this study and may need additional follow-up experiments.

This study highlights the feasibility of adopting measurements from protein-derived amino acids, which are stable to analyse using GC-MS, to undertake steady-state MFA using various 13C tracers to define the metabolic phenotypes of A549 cells better.

In perturbed conditions like cancer and infectious diseases, metabolic hyperactivity with increased protein catabolism and negative nitrogen balance is commonly observed. So, understanding the dynamic changes in amino acids and primarily the pathways that contribute to their synthesis or their precursor molecules could be useful to develop nutritional supplement for disease management, including improving its outcome. Pathogens, like *Mycobacterium leprae*, uses host glucose pool to synthesize the majority of amino acids, and these related pathways are useful to target for anti-leprosy drug development [10]. Exogenous alanine in T-cells is not used in metabolism but utilised for protein synthesis and its normal activation [11]. So, targeting the evolutionarily conserved phenomenon of protein synthesis by modulating amino acid metabolism could favourably influence the host immune responses in cancer, infection and autoimmunity.

In summary, the workflow adopted to monitor the fate of 13C glucose analysis and deriving average 13C of labelled metabolite provides an optimised tool that could be useful in metabolic phenotypes of cells exposed to perturbed conditions, including viral and bacterial infection. It may also be helpful to understand the anterograde and retrograde signalling in mitochondria that provides cues for uncontrolled cell proliferation.

## 4. Materials and Methods

### 4.1 Cell Culture

Type-II alveolar epithelial adenocarcinoma cell line, A549, was procured from National Centre for Cell Science (NCCS), Pune (India). Cells (7^th^ passage) were cultured in six-well plates (0.5 million cells/well) containing 2.0 ml DMEM-F12 (Catalogue No. L0091-500, Biowest, USA) and 10% heat-inactivated fetal bovine serum (FBS, Catalogue No. 10270-106, South American Origin, GIBCO) with 17.5 mM U-^13^C_6_ glucose (Catalogue No. GLC-082, Omicron, USA) in triplicates at humidified 37 °C with 5% CO_2_ for 8 hours for adherence and treated as the first time point.

### 4.2 Growth curve and viability of A549 cells across different time-points

A549 cells, were cultured in 6-well plate and harvested after every 12 hours (upto 60 hours) for cell counting and Trypan Blue viability assay.

### 4.3 Metabolites extraction

At every 12 hours up to 60 hours, cell culture filtrate (ccf) was collected and cells were washed with saline solution (0.9% NaCl) and chilled methanol (800 μl) was used for quenching followed by milli-Q water (800 μl). Cells were scraped, collected in centrifuge tubes and after adding chloroform (1.6 ml) were vortexed at 4 °C for 30 min. These samples were centrifuged at 10,000 × g for 10 min at 4 °C, and the aqueous phase samples were harvested and stored at -20 °C until further analysis.

### 4.4 Pre-processing of samples for GC-MS

Metabolite extracts from ccf (60 μl) and methanolic extract (300 μl) samples, with internal standard (10 μl of 2 mg/ml Isoniazid) were dried using a speed vacuum evaporator. Metabolites were derivatised by incubating with 40 μl Methoxyamine hydrochloride (MOX) in pyridine followed by 80 μl N-Methyl-N-(trimethylsilyl) trifluoroacetamide (MSTFA) at 37 °C for 2 hours and 30 min respectively. The junction layer of the aqueous (methanolic) and organic (chloroform) phases comprising of proteins was selected for amino acid analysis. Protein samples were hydrolysed and derivatised following earlier published method [12]. All the processed samples were centrifuged (10 min, 16,100 × g at RT), and the supernatant was taken for GC-MS analysis.

### 4.5 GC-MS Data acquisition

All samples were run in a single batch. Derivatized samples (1 μl) were injected into the GC column (Restek, RTX-5, 30 m x 250 μm x 0.25 μm)) in a purged splitless mode. Helium was used as the carrier gas (0.6 ml/min), with a solvent delay of 12 min and the oven program was set at 50 °C for 5 min, a 5 °C/min ramp to 200 °C, held for 10 min, a 20 °C/min ramp to 300 °C and held for 4 min. For TBDMS derivatized samples, Helium gas was used as carrier gas (1.3 ml/min) with a different oven programme starting with 120 °C for 5 min, 4 °C/min ramp to 270 °C, held for 3 min, a 20 °C/min ramp to 320 °C, held for 1 min. The EI ion source temperature was set at 230 °C, quadrupole temperature set at 150 °C and data was acquired in full scan mode from m/z 50 to m/z 600. Agilent Chemstation (G1701DA GC/MSD) was used during data acquisition and for peak integration. Metabolite identification was confirmed by running commercial standards and National Institute of Standards and Technology (NIST) database search.

### 4.6 Data analysis

GC-MS spectra were baseline corrected using MetAlign [13] to estimate the accurate mass isotopomer distribution (MIDs) for all amino acids. The incorporated natural abundance of 13C isotope contributes to the measured MIDs. For accurate estimation of mass isotopomer abundances, the correction of MIDs was performed using Isocor software [14]. The finally corrected MIDs were then used to calculate average ^13^C abundance in each amino acid fragment [15].

### 4.7 Statistical Analysis

Error bars represent standard deviation between experimental replicates. The number of replicates is indicated in figure legends. Plots were prepared using GraphPad Prism version 8.0.1. Figures were created with BioRender.com

## Author Contributions

Conceptualization, S.A. and R.K.N.; methodology, S.A. and R.K.N.; software, S.K.M..; validation, S.A., S.K.M., J.T. and R.K.N.; formal analysis, S.A., J.T., M.S. and S.K.M.; investigation, S.A.; resources, R.K.N. and S.K.M.; data curation, S.A.; writing—original draft preparation, S.A., S.K.M. and R.K.N.; writing—review and editing, S.A., J.T., M.S., S.K.M. and R.K.N.; visualization, S.A.; supervision, R.K.N.; project administration, R.K.N.; funding acquisition, R.K.N. and S.K.M. All authors have read and agreed to the published version of the manuscript.

## Funding

The study was supported by CORE funding from International Centre for Genetic Engineering and Biotechnology New Delhi to R.K.N. and SKM acknowledges the seed grant support of Indian Institute of Technology Mandi. Research Fellowship from Department of Biotechnology, Government of India to S.A. and PhD fellowship from Ministry of Human Resource Development, Government of India to J.T. and M.S. are acknowledged.

## Data Availability Statement

The data presented in this study are available in Mendeley Data at <u>10.17632/gc3ss3s4jx.1</u>.

## Acknowledgments

We thank the Translational Health Laboratory members for technical help and comments on the manuscript.

## Conflicts of Interest

The authors declare no conflict of interest.

## Notes

http://dx.doi.org/10.17632/gc3ss3s4jx.1

### Competing Interest Statement

The authors have declared no competing interest.

### Summary of Updates

We have revised the abstract, restructured the manuscript by moving the detailed methodology from supplementary file to the main text, Figure 1 is presented as two separate figures and updated in the revised manuscript with a discrete discussion section.

## References

1. Kurmi, K.; Haigis, M. C. Nitrogen metabolism in cancer and immunity. Trends Cell Biol. 2020, 30, 408–424.

2. Kelly, B.; Pearce, E. L. Amino assets: how amino acids support immunity. Cell Metab. 2020, 32, 154–175.

3. Jain, M.; Nilsson, R.; Sharma, S.; Madhusudhan, N.; Kitami, T.; Souza, A. L.; Kafri, R., Kirschner; M. W., Clish; C. B.; Mootha, V.K. Metabolite profiling identifies a key role for glycine in rapid cancer cell proliferation. Science. 2012, 336, 1040–1044.

4. Hosios, A. M.; Hecht, V. C.; Danai, L. V.; Johnson, M. O.; Rathmell, J. C.; Steinhauser, M. L.; Manalis, S. R.; Vander Heiden, M. G. Amino acids rather than glucose account for the majority of cell mass in proliferating mammalian cells. Dev. Cell. 2016, 36, 540–549.

5. Cambridge, S. B.; Gnad, F.; Nguyen, C.; Bermejo, J. L.; Krüger, M.; Mann, M. Systems-wide proteomic analysis in mammalian cells reveals conserved, functional protein turnover. J. Proteome Res. 2011, 10, 5275–5284.

6. Chen, P. H.; Cai, L.; Huffman, K.; Yang, C.; Kim, J.; Faubert, B.; Boroughs, L.; Ko, B.; Sudderth, J.; McMillan, E. A.; et al. Metabolic diversity in human non-small cell lung cancer cells. Mol. Cell. 2019, 76, 838–851.e5.

7. Liu, S.; Dai, Z.; Cooper, D. E.; Kirsch, D. G.; Locasale, J. W. Quantitative analysis of the physiological contributions of glucose to the TCA cycle. Cell Metab. 2020, 32, 619– 628.e21.

8. Hui, S.; Cowan, A. J.; Zeng, X.; Yang, L.; TeSlaa, T.; Li, X.; Bartman, C.; Zhang, Z.; Jang, C.; Wang, L.; et al. Quantitative fluxomics of circulating metabolites. Cell Metab. 2020, 32, 676–688.e4.

9. Hu, X.; Chao, M.; Wu, H. Central role of lactate and proton in cancer cell resistance to glucose deprivation and its clinical translation. Signal Transduct. Target Ther. 2017, 2, 16047.

10. Borah, K.; Girardi, K.; Mendum, T. A.; Lery, L.; Beste, D.; Lara, F. A.; Pessolani, M.; McFadden, J. Intracellular Mycobacterium leprae utilizes host glucose as a carbon source in Schwann cells. MBio. 2019, 10, e02351–19.

11. Ron-Harel, N.; Ghergurovich, J. M.; Notarangelo, G.; LaFleur, M. W.; Tsubosaka, Y.; Sharpe, A. H.; Rabinowitz, J. D.; Haigis, M. C. T Cell activation depends on extracellular alanine. Cell Rep. 2019, 28, 3011–3021.e4.

12. Shree, M.; Masakapalli, SK. Intracellular fate of universally labelled ^13^C isotopic tracers of glucose and xylose in central metabolic pathways of Xanthomonas oryzae. Metabolites 2018, 8, 66.

13. Lommen, A.; Kools, HJ. MetAlign 3.0: performance enhancement by efficient use of advances in computer hardware. Metabolomics 2012, 8, 719–726.

14. Millard, P., Letisse, F., Sokol, S., Portais, JC. IsoCor: correcting MS data in isotope labeling experiments. Bioinformatics 2012, 28, 1294–1296.

15. Masakapalli, SK.; Kruger, NJ.; Ratcliffe, RG. The metabolic flux phenotype of heterotrophic Arabidopsis cells reveals a complex response to changes in nitrogen supply. Plant J. 2013, 74, 569–582.

